# The accumulation of chloroplast small RNAs in unicellular alga *Chlamydomonas reinhardtii* is affected by nitrogen deprivation

**DOI:** 10.1101/839092

**Authors:** Suhaimi Che-Ani, Ghows Azzam, Nazalan Najimudin

**Author notes:** Penang, Malaysia 11800. <u>Corresponding authors:</u> Nazalan Najimudin, School of Biological Sciences, Universiti Sains Malaysia, 11800 Minden, Penang, Malaysia.

## Abstract

Small RNAs generated from the chloroplast genome may play a role in gene regulation. Given that chloroplast function is affected by nitrogen deprivation, there is yet an attempt to link chloroplast small RNAs to this stress condition. This study aims to determine the response of chloroplast small RNAs under nitrogen deprivation and their putative mode of action. A comparative transcriptomic approach was carried out to analyze the differential accumulation of chloroplast small RNAs from *Chlamydomonas reinhardtii* cells grown in nitrogen-deprived versus nitrogen-based medium. A total of 101 chloroplast small RNA candidates were successfully annotated. Growth in nitrogen-deprived medium revealed 17 significantly upregulated and 12 downregulated chloroplast small RNAs. These chloroplast small RNAs originated from different genomic locations such as untranslated, intergenic or antisense regions as well as the ends of tRNA and rRNA genes. The differentially accumulated csRNAs from 3’-untranslated regions were all upregulated. In contrast, the csRNAs from the ends of tRNA and rRNA genes were all downregulated during nitrogen deprivation. Fluctuations of the chloroplast small RNA levels indicated their importance in the chloroplasts during changes in nitrogen levels. The primary sequences of three selected chloroplast small RNA were found to be conserved in the chloroplast genomes of a few microalgae, again reflecting their functional importance. The findings from this study provided new insights into the involvement of non-coding RNAs in chloroplast during metabolic stress.

## INTRODUCTION

Non-coding RNAs (ncRNAs) are RNA molecules that are not translated into polypeptides or proteins. Besides ribosomal RNA and transfer RNA, ncRNAs also include bacterial small RNA, microRNAs, small interfering RNAs, antisense RNAs, long non-coding RNA, etc. Their sizes range from 18 to 500 nt long. They have been increasingly recognized as important factors in regulating gene expression at the transcriptional and post-transcriptional level (Kawaji and Hayashizaki 2008; Ghildiyal and Zamore 2009; Cech and Steitz 2014). These multifunctional molecules have also been reported to be involved in DNA replication, splicing, translation, and genome defense (as reviewed in Kim 2005; Mattick and Makunin 2006). Non-coding RNAs are found in a broad spectrum of organisms and their roles have been established in several model organisms such as *Escherichia coli*, *Caenorhabditis elegans*, *Drosophila melanogaster*, *Arabidopsis thaliana*, mice and human (Bartell 2004; Axtell *et al*. 2006; Bonnet *et al*. 2006; Yigit *et al*. 2006). They are produced at different development stages or in response to various external stimuli suggesting that they are functionally involved in numerous physiological processes (as reviewed in Gottesman *et al*. 2004; Jonathan *et al*. 2007).

In photosynthetic organisms, the location of ncRNAs plays an important part in their regulatory roles (Leung and Sharp 2006). Molecules of ncRNAs were shown to be present not only in the nuclear-cytosolic compartment but also in the chloroplasts of *Chlamydomonas reinhardtii* (Goldshmidt-Clermont *et al*. 1991; Cavaiuolo *et al*. 2017), *Arabidopsis thaliana* (Lung *et al*. 2006; Georg *et al*. 2010; Hotto *et al*. 2011; Zghidi-Abouzid *et al*. 2011; Cognat *et al*. 2017; Ruwe *et al*., 2016), *Nicotiana tabacum* (Hotto *et al*. 2010), *Brassica rapa* (Wang *et al*. 2011), *Cucumis melo* (Gonzalez-Ibeas *et al*. 2011), *Hardeum vulgare* (Zhelyazkova *et al*. 2012; Hackenberg *et al*. 2015), *Salvia miltiorrhiz* (Chen *et al*. 2014) and *Silene noctiflora* (Wu *et al*. 2015). Small RNA libraries from purified chloroplasts confirmed the existence of chloroplast-encoded ncRNAs which were later were abbreviated as <u>c</u>hloroplast <u>s</u>mall <u>RNAs</u> (csRNAs) (Wang *et al*. 2011). Accumulation of csRNAs in chloroplast could be due to relaxed initiation and inefficient termination of transcription resulting in the generation of full transcripts from both chloroplast DNA strands. This led to the transcription of all regions including non-coding regions, which consequentially enabled the production of csRNAs (Hotto *et al*. 2012; Shi *et al*. 2016). Several investigations were made to understand the modes of actions and physiological roles of csRNAs at different developmental stages of growth and environmental conditions, especially in transcriptional and post-transcriptional regulation (Hotto *et al*. 2011; Wang *et al*. 2011; Ruwe and Schmitz-Linneweber 2012; Hackenberg *et al*. 2015; Ruwe *et al*. 2016).

N is an integral component of life and is required for growth and development. In photosynthetic eukaryotes, N deprivation has been shown to adversely affect their general transcriptional and metabolomic profiles (Peltier and Schmidt 1991; Miller *et al*. 2010; Msanne *et al*. 2012). The effect also included the regulation of photosynthetic structure and function (Schmollinger *et al*. 2014; Juergens *et al*. 2015). Photosynthetic functions involving photosystems I and II decreased along with the lipid levels in the thylakoid membrane although the latter appeared to be preserved. As chloroplast is the factory of photosynthesis, the expression of chloroplast genes decreased during N deprivation (Plumley and Schmidt 1989). These observations suggested that N availability is a very significant variable to be explored, especially on their effect on chloroplast function. Moreover, there is limited information on the response of csRNAs during N deprivation.

In this study, a comparative transcriptomic analysis was performed to investigate the relative levels of csRNAs in response to N deprivation in the model unicellular algae *Chlamydomonas reinhardtii*. We used the deep next-generation sequencing approach to identify changes in the csRNAs level in N-deprived (ND) and N-based (NB) growth media. Several csRNAs that were significantly differentially accumulated were further analyzed based on their location on the chloroplast genome. The possibility of these csRNAs to target protein-coding genes during N deprivation is also discussed. Thereafter, we determined the sequence of the full transcripts of selected csRNAs to observe their secondary structures and analyze their sequence conservation.

## MATERIALS and METHODS

### *Chlamydomonas* strain and culture conditions

*Chlamydomonas reinhardtii* strain CC-503 cw92 mt^+^ was obtained from Chlamydomonas Resource Centre, University of Minnesota (http://chlamycollection.org/). *Chlamydomonas* cells were grown mixotrophically in Tris-Acetate-phosphate (TAP) liquid medium at 25°C shaken at 100rpm under constant light. Cell growth was measured by determining the optical density at 750nm using the Beckman-Coulter DU 800 Spectrophotometer. For the comparative transcriptomic study, the strain was cultured in 100ml TAP for 48h until the early log phase with an OD of approximately 0.2. The cells were centrifuged, and the resultant pellet was transferred to a fresh volume of 100ml TAP-N (TAP medium lacking ammonium chloride as N source) to initiate N deprivation condition. Growth was continued for another 24 h before the cells were harvested for the RNA extraction procedure (Juergen *et al*. 2015).

### RNA preparation and small RNA Sequencing

Two independent biological replicates for each condition were used for the small RNA library construction using the Solexa high-throughput sequencing platform. After 24 hours of growth, total cells were separately collected from the TAP and TAP-N media by centrifugation at 12000g for 10min and washed with PBS buffer. Total RNA samples from approximately 5 x 10^8^ cells were extracted using mirVana™ miRNA Isolation Kit (Ambion). RNA concentration was measured by measuring absorbance at 260nm using the Eppendorf Biospectrometer. Genomic DNA was removed using the TURBO DNA-free kit (Ambion). Total RNA samples were sent to BGI Tech. Solutions Co (Hong Kong) for RNA quality assessment and small RNA sequencing. The libraries were prepared and sequenced on Illumina-HiSeq25000 according to the manufacturer’s protocol. The “clean reads” were provided by the manufacturer after the removal of low-quality reads and adaptor sequences.

### Bioinformatics analysis of chloroplast mapped small RNAs

The clean reads were mapped to the *C. reinhardtii* chloroplast (GenBank accession number: NC_005353), nuclear (JGI v4.0) and mitochondrial genomes (Genbank accession number: NC_001638) using the software *butter* (bowtie utilizing iterative placement of repetitive small RNAs) software with no mismatch (Axtell 2014). Reads that mapped to the intergenic and antisense regions of the chloroplast genome were pooled from all datasets and assembled to form clusters using the *bedtools merge* software. For clusters that originated from inverted repeat regions, only clusters that came from inverted repeat A were included in the analysis to avoid redundancy. The coverage in the chloroplast genome generated by the clusters was viewed using Integrative Genomics Viewer (IGV) browser (Broad Institute). For differential analysis, the counts of csRNAs were defined as the number of reads mapped to the clusters. The counts for each dataset were normalized and statistically validated using the software *DESeq* under default parameter for comparative analysis (Ander and Huber 2010).

### Quantitative validation of csRNA levels using real-time quantitative PCR assay

The presence and relative accumulation of csRNAs were validated using real-time quantitative PCR (qRT-PCR) assay. The method used was according to the protocol involving polyadenylation of small RNAs (Ro and Yan 2010). Small RNAs were extracted from DNAse-treated total RNA using the modified MirVana (Ambion) protocol without the columns. Subsequently, 2ug of small RNA samples were added with poly(A) tails at their 3’ ends using the Poly(A) Tailing Kit for Polyadenylation of RNA (Applied Biosystems). Small RNA samples from this reaction were purified again using the phenol: chloroform and ethanol precipitation method. For cDNA synthesis, 2ug of tailed small RNAs were reverse transcribed using the SuperScript™ First-Strand Synthesis System (Invitrogen, USA) and its miRTQ adapter primer containing oligo dTs. The qRT-PCR assay was performed using polyadenylated cDNAs as a template, core sequence of ccsRNAs as the forward primers and RTQ-UNIr as the reverse primer. Genes coding for chloroplast 3S rRNA (*rrn3*), U4 small nucleolar RNA (U4 snRNA) and U6 small nucleolar RNA (U6 snRNA) were used as references (Shu and Hu 2012). The primers (IDT DNA Technologies) used are listed in Table S2. Every qRT-PCR reaction was carried out in a 10μL volume containing: 5 μL *SsoAdvanced Universal SYBR Green Supermix* (Applied Biosystems), 1 uL template, 1 uL forward primer, 1 uL reverse primer and 3 uL nuclease-free H_2_O. qRT-PCR was performed in a CFX96 real-time PCR detection system (Bio-Rad Laboratories, USA) for 30 s at 95°C, followed by 40 runs of 10 s at 95°C, 10 s at 55°C and 20 s at 72°C. For each reaction, a ‘No Template’ control (NTC) was included. Each reaction was technically triplicated. The PCR products were ligated into pGEMT-Easy and sequenced using a Sanger-based method for sequence confirmation.

### Analysis of qRT-PCR dataset

The specificity of amplification using designated primers was evaluated based on melting curve analysis and product sizes. A standard curve was implemented for each primer set using serial dilutions of DNA templates to test for the amplification efficiency. Data analysis was performed with the CFX96 real-time detection system (Bio-Rad) with automatic Cq and baseline setting using reference genes as the internal controls. Normalized relative csRNA accumulations were calculated using the *Vandosampele* method (Vandosampele *et al*. 2002).

### Determination of 5’ and 3’ ends of csRNA primary transcripts by intramolecular circularization

The termini of csRNA primary transcripts were determined by modified RT-PCR amplification of 5’ and 3’-circularised RNA molecules (Slomovic and Schuster 2013). The circularization of DNase treated total RNAs was performed by intramolecular ligation of 5’ and 3’ ends using 40U of T4 RNA ligase (New England Biolabs) according to the manufacturer’s protocol. The reaction mixture was purified using the phenol: chloroform and ethanol precipitation method (Ro and Yan 2010). For cDNA synthesis, 2ug of circular RNA were reverse transcribed using SuperScript™ First-Strand Synthesis System (Invitrogen, USA) and a specific reverse primer that bound to the core sequence of csRNAs. The single-stranded cDNAs were subsequently amplified using csRNA specific primers that were derived from the sequences that flanked each core sequence as listed in Table S3. The Intron (NHK Biosciences) DNA purification kit was used to purify the PCR products which were subsequently ligated into pGEMT-Easy for the DNA sequencing process.

### Bioinformatics analysis of primary transcript of csRNAs

The secondary structures of the primary csRNA transcripts with the lowest minimum free energy were predicted using the *RNAFold* program under default parameter (Hofacker 2003). To search for sequence similarity in different organisms, the primary csRNA sequences were subjected to BLASTN analysis under both high and low stringency parameters. Similar sequences that were found in photosynthetic organisms were pooled for phylogenetic analysis using the program *BLAST Tree View*.

### Data Availability Statement

*Chlamydomonas reinhardtii* strain is available at Chlamydomonas Resource Centre, University of Minnesota. Supplementary File contains Table S1, S2, S3, and S4 (The file was uploaded to the GSA Figshare portal). The authors affirm that all data necessary for confirming the conclusions of the article are present within the article, figures, and supplemental materials. The gene expression data will be uploaded to the Gene Expression Omnibus (GEO) repository for an accession number.

## RESULTS

### Read mapping analysis against the *C. reinhardtii* chloroplast genome

Chloroplast functions have been known to be regulated by the availability of N. Thus understanding the differential presence of chloroplast small RNA species of cells grown in different N availability was crucial. Equal numbers of algal cells were harvested 24 h after N deprivation. This span of time was chosen based on the previous time-course study of differential expression levels of transcripts and proteins of photosynthesis (Juergens *et al*. 2015). Two independent biological replicates for each condition were generated (Figure 1A). Each of the four small RNA libraries constructed for *C. reinhardtii* cells under N-based (NB1 and NB2 libraries) and N-deprived (ND1 and ND2 libraries) conditions produced around 5 million high-quality sequence reads of 18 to 44 nt in length (Table 1A).

**Figure 1:**
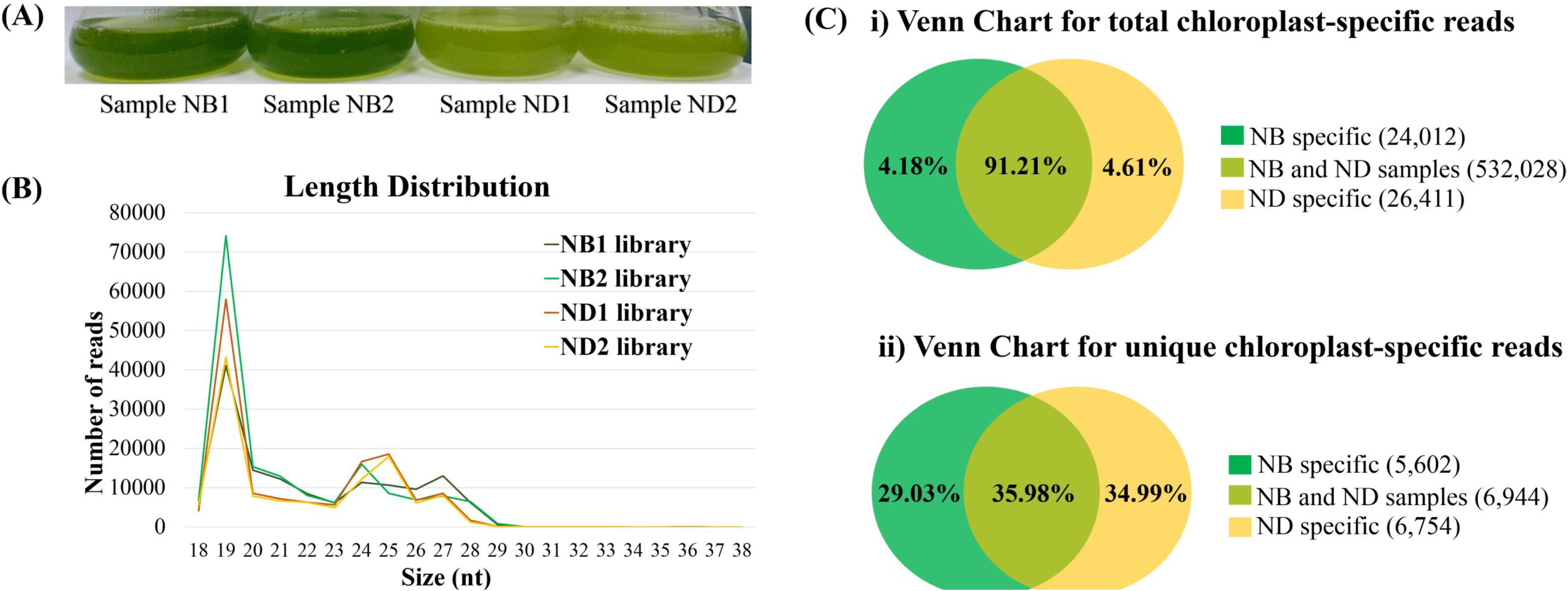
The summary of library construction and *in silico* analysis of chloroplast small RNAs. (A) The samples of *C. reinhardtii* cells under N-based (NB1 and NB2) and N-deprived (ND1 and ND2) used for small RNA library construction. (B) The size distribution of chloroplast-specific reads in NB1, NB2, ND1, and ND2 libraries. (C)Venn Charts to demonstrate the common and specific sequences of (i) total and (ii) unique chloroplast-specific reads. [The Venn charts were presented in percentage value and the figures were not drawn into scale. Number of each common and specific reads for each condition were shown inside brackets in legends.]

**Table 1.** Analysis of production and alignment of small RNA reads.

| Samples | NB1 | NB2 | ND1 | ND2 |
| --- | --- | --- | --- | --- |
| A. Total reads generated from high-throughput sequencing | 5,363,812 | 5,332,446 | 5,830,736 | 5,632,690 |
| B. Chloroplast Genome Alignment |  |  |  |  |
| I. Total Chloroplast-mapped reads | 141,090 | 170,666 | 142,762 | 120,565 |
| II. Number of chloroplast-mapped reads that overlapped with nuclear and mitochondrial genome | 392 | 456 | 416 | 368 |
| C. Chloroplast-specific reads |  |  |  |  |
| I. Total reads | 140,698 | 170,210 | 142,346 | 120,197 |
| II. Percentage from whole population* | 2.62 % | 3.12 % | 2.44 % | 2.13 % |
| III. Unique reads | 5,258 | 5,620 | 6,433 | 6,058 |

Chloroplast-mapped reads were obtained by aligning total reads to the chloroplast genome of *C. reinhardtii* with a perfect match using the software *butter*. This software eliminated the biasness of random mapping of multi mapped reads since these reads are placed to regions of confidently high density. Some of the chloroplast-mapped reads additionally mapped into the nuclear and mitochondrial genomes of *C. reinhardtii*. The incidence of multi mapped reads were low and are shown in Table 1B. Multimapped reads were omitted to obtain chloroplast-specific reads. The replicates of N-based condition produced 140,698 (NB1) and 170,210 (NB2) chloroplast-specific reads while those of N-deprived condition produced 142,346 (ND1) and 120,197 (ND2) chloroplast-specific reads. The reads from all the libraries exhibited wide variations in length, from 18 to 38 nt with 19 nt was the most abundance reads (Figure 1B). The chloroplast alignment represented around 2-3 % of the whole population of reads (Table1C). Identical sequences from the chloroplast-specific reads were further grouped into single reads and unique reads were subsequently determined from these (Table 1C). As much as 36% of unique reads were found to be common in NB and ND conditions, while 29% and 35% of unique reads were NB specific and ND specific, respectively (Figure 1C).

### The annotation of chloroplast small RNAs in *Chlamydomonas reinhardtii*

Non-coding RNA comes in several groups in eukaryotes with transfer RNA (tRNA) and ribosomal RNA (rRNA) being the most prominent. Besides these key groups, a number of other noncoding RNA classes also exist in eukaryotic cells. Reads that mapped to the non-coding regions are prime targets of chloroplast small RNA analysis. These exclude reads from sense open reading frame, tRNA and rRNA. A total of 1337 clusters were formed when the reads were assembled. These clusters were filtered according to the following criteria to increase the confidence of the detection level. Firstly, only the clusters representing more than 50 reads were retained. Secondly, only clusters demonstrating a narrow peak of sequence density were considered (as recommended by Ruwe and Schmitz-Linneweber 2016). The coverages of clusters were viewed in Integrated Genomic View (IGV) browser to exclude those having broad regions of sequence density. A set of 101 clusters that fulfilled the criteria were identified, which were henceforth called Cre-csRNA (*<u>C</u>hlamydomonas <u>reinhardtii csRNA</u>*). The most abundant unique read defined the core sequence of each csRNA as shown by an example (Cre-csRNA_022) in Figure 2.

**Figure 2:**
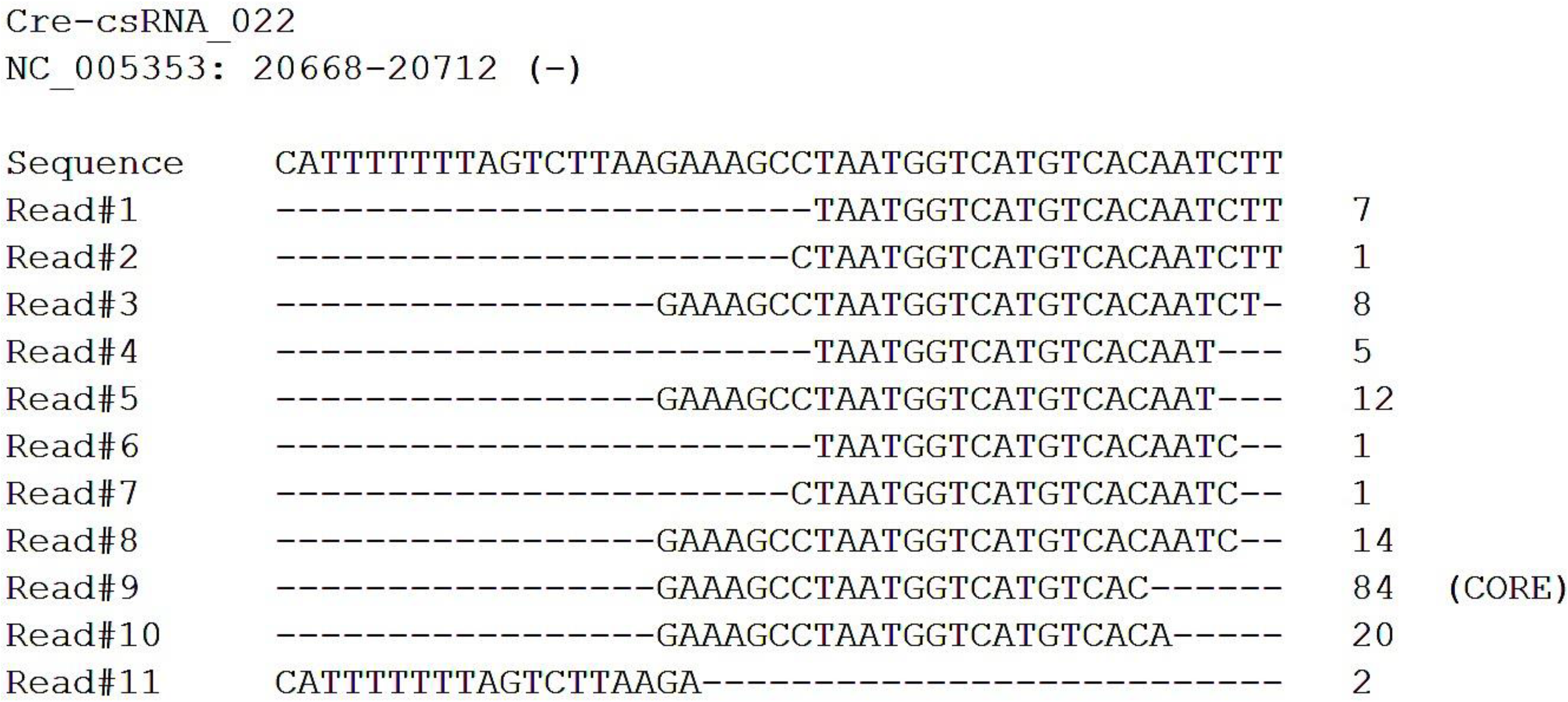
An example of the identification of the core sequence in csRNAs. The most abundant unique read in Cre-csRNA_022 is Read#8 and the sequence of this read is defined as core sequence for Cre-csRNA_022.

### Differential accumulation of csRNAs in N-deprived versus N-based conditions

The counts of aligned reads into 101 csRNAs for each dataset (NB1, NB2, ND1, and ND2) were normalized and statistically validated using *DESeq* software. A total of 32 csRNAs were discovered to be differentially accumulated significantly upon N deprivation using an adjusted p-value of < 0.05. A total of 17 csRNAs were found to be upregulated (log_2_ fold change > 1) and 12 csRNAs were downregulated (log_2_ fold change < −1) as listed in Tables 2 and 3. Three csRNAs were found to be unique and only present in the NB dataset as listed in Table 4. The core sequences of each N-responsive csRNA were summarized in Table S1.

**Table 2.**
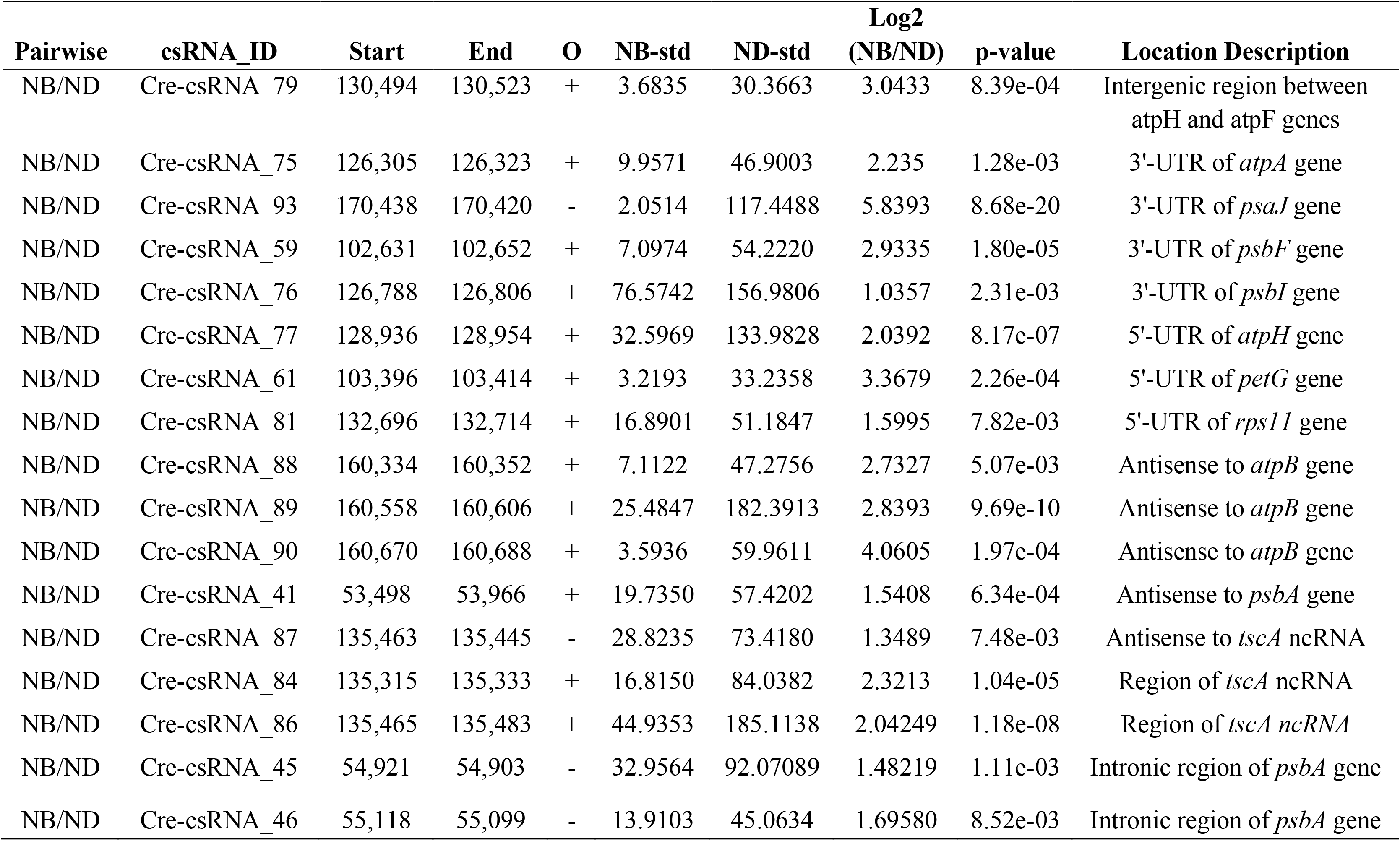
List of upregulated csRNAs (log2 Fold Change > 1; p-value < 0.05)

**Table 3.** List of downregulated csRNAs (log2 Fold Change < −1; p-value < 0.05)

| Pairwise | csRNA_ID | Start | End | O | NB-std | ND-std | Log2 | p-value | Location Description |
| --- | --- | --- | --- | --- | --- | --- | --- | --- | --- |
|  |  |  |  |  |  |  | (NB/ND) |  |  |
| NB/ND | Cre-csRNA_98 | 191,999 | 191,217 | + | 52.6464 | 6.9395 | -2.9234 | 4.87e-05 | Intergenic region between *trnN* and *rpoC1b* genes |
| NB/ND | Cre-csRNA_17 | 14,990 | 14,972 | - | 1211.3179 | 472.5245 | -1.3581 | 1.65e-05 | 3' end of *trnC* tRNA gene |
| NB/ND | Cre-csRNA_96 | 190,887 | 190,905 | + | 425.1673 | 180.8035 | -1.2336 | 7.96e-05 | 3' end of *trnN* tRNA gene |
| NB/ND | Cre-csRNA_18 | 15,152 | 15,134 | - | 117.1518 | 52.8677 | -1.1479 | 4.57e-03 | 3' end of *trnT* tRNA gene |
| NB/ND | Cre-csRNA_65 | 118,429 | 118,453 | + | 59.7734 | 23.5877 | -1.3415 | 1.60e-02 | 3' end *trnQ* tRNA gene |
| NB/ND | Cre-csRNA_30 | 41,750 | 41,772 | + | 202.1256 | 25.2170 | -3.0028 | 4.18e-14 | 5' end of *rrn7* rRNA gene |
| NB/ND | Cre-csRNA_82 | 133,768 | 133,792 | + | 60.4925 | 24.0772 | -1.3291 | 1.80e-02 | 5' end of *trnK* tRNA gene |
| NB/ND | Cre-csRNA_91 | 161,954 | 161,931 | - | 175.1589 | 70.7093 | -1.3087 | 1.90e-04 | 5'-UTR of *atpB* gene |
| NB/ND | Cre-csRNA_22 | 20,695 | 20,674 | - | 54.6979 | 10.9558 | -2.3198 | 5.20e-04 | 5'-UTR of *petB* gene |
| NB/ND | Cre-csRNA_62 | 107,987 | 107,966 | - | 90.1091 | 16.4268 | -2.4556 | 3.90e-04 | Antisense to 5'-UTR of *rpoC2* gene |
| NB/ND | Cre-csRNA_70 | 123,567 | 123,588 | + | 37.9278 | 3.1982 | -3.5679 | 2.55e-03 | Antisense to *rbcL* gene |
| NB/ND | Cre-csRNA_71 | 123,833 | 123,855 | + | 22.5948 | 4.5525 | -2.3113 | 2.27e-02 | Antisense to *rbcL* gene |

**Table 4.** List of NB-specific csRNAs (baseMeanND = 0; p-value < 0.05)

| Pairwise | csRNA_ID | Start | End | O | NB-std | ND-std | Log2<br>(NB/ND) | p-value | Location Description |
| --- | --- | --- | --- | --- | --- | --- | --- | --- | --- |
| NB/ND | Cre-csRNA_10 | 7,439 | 7,457 | + | 28.2102 | 0 | NA | 1.27e-02 | 5'-UTR of *chlB* gene |
| NB/ND | Cre-csRNA_25 | 28,733 | 28,716 | - | 23.7178 | 0 | NA | 6.71e-05 | Antisense to 5'-UTR of<br>*rpl14* gene |
| NB/ND | Cre-csRNA_66 | 120,180 | 120,159 | - | 51.7630 | 0 | NA | 8.25e-10 | Antisense to *psaB* gene |

The locations of each N-responsive csRNAs on the chloroplast genome were investigated and stated in Tables 2,3 and 4. The assignment of the positions in terms of 5’ and 3’ untranslated regions (UTR) of chloroplast protein-coding genes was based on Cavaiuolo *et al*. 2017. Twelve out of the 32 differentially accumulated csRNAs were found to be located at either the 5’ and 3’ UTR of protein-coding genes. Among csRNAs located at 5’-UTR of protein-coding genes, 3 were upregulated and 3 were downregulated. Interestingly, 2 of these were only present in the NB dataset. Three csRNAs located at 3’-UTR of protein-coding genes were differentially accumulated and they were all upregulated under N-deprived condition. Apart from those that are positioned at UTRs of protein-coding genes, several csRNAs were also found in the ends of chloroplast tRNA and rRNA genes. Five were located at 5’ or 3’ end of tRNA genes and one was positioned at 5’ end of an rRNA gene. They were all downregulated. The number of N-responsive csRNAs located from the non-transcribed intergenic region was very low. Only two were found and one was upregulated while the other was downregulated.

Differentially accumulated csRNAs that were antisense to several protein-coding genes were also uncovered. Three were antisense to the *atpB* gene and one was antisense to the *psbA* gene, and they were all upregulated. Remarkably, two downregulated csRNAs were antisense to the *rbcL* gene. The intronic region also contributed to the generation of N-responsive csRNAs. The chloroplast genome has two homologs of the *psbA* gene and both have introns. Two csRNAs were revealed and both were upregulated. The *Chlamydomonas* chloroplast genome also has a non-coding RNA gene that is involved in trans-splicing reaction. This gene is called *tscA* and three upregulated csRNAs were mapped to it.

### Validation of small RNA-Seq using RT-qPCR

All the 32 differentially accumulated csRNAs were subjected to pre-screening to verify their presence by reverse transcriptase-PCR. The expression of these csRNAs were further measured by using RT-qPCR. Eight passed the requirements for RT-qPCR analysis, while the rest failed to establish an acceptable standard curve. The validated transcripts included four upregulated csRNAs (Cre-csRNA_45, Cre-csRNA_61, Cre-csRNA_75 and Cre-csRNA_79) and four downregulated csRNAs (Cre-csRNA_17, Cre-csRNA_30, Cre-csRNA_62 and Cre-csRNA_91). The normalized relative expression is summarized in Figure 3A and 3B. The positive correlation validated the result of the differential expression of the transcriptomic analysis. Unfortunately, attempts to validate the three NB-specific csRNAs were not performed because they were non-detectable at the pre-screening FRT-PCR stage.

**Figure 3:**
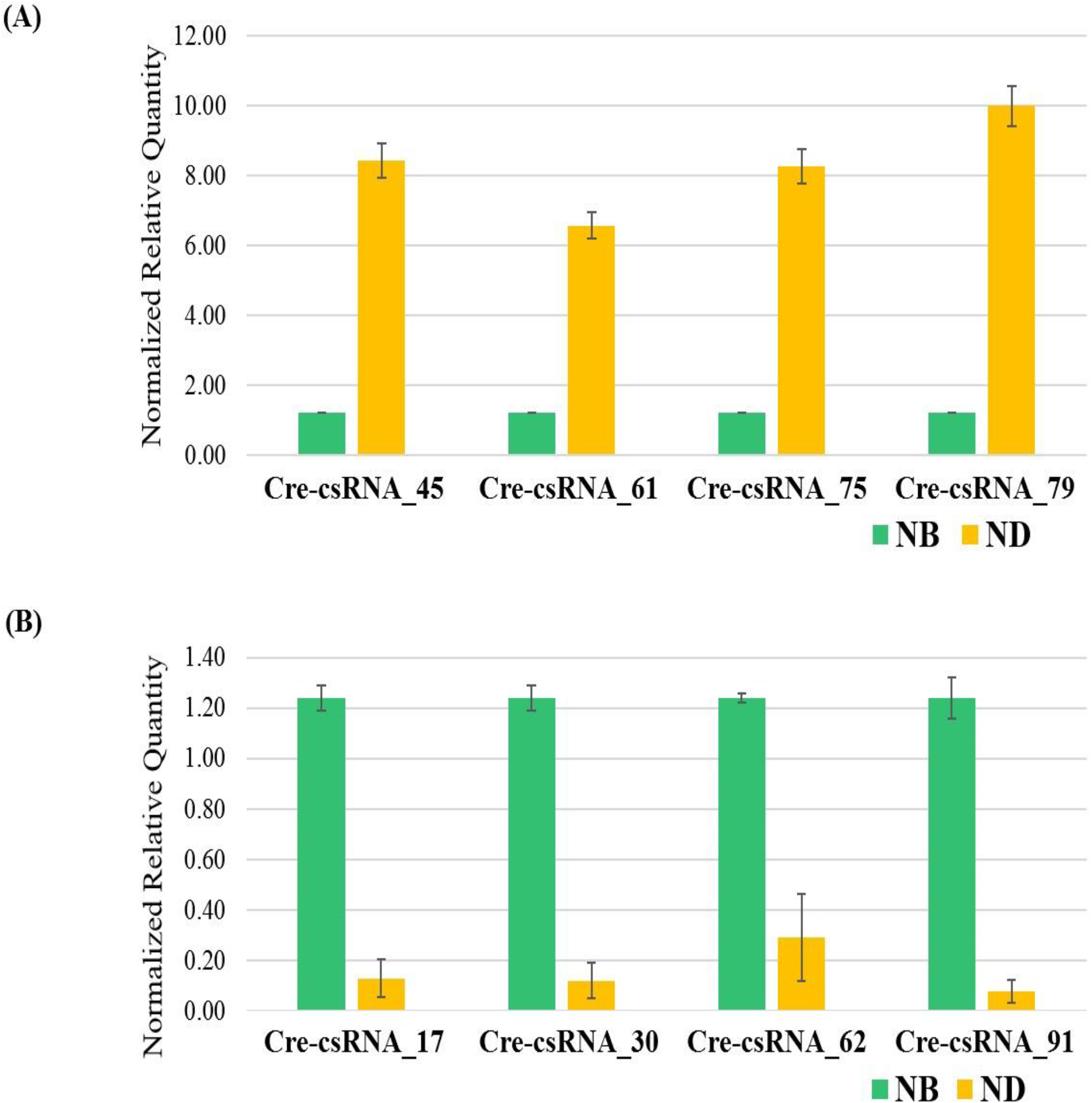
Quantitative RT-PCR analysis of N-responsive csRNAs. Normalized fold changes for the expression of four upregulated (A) and four downregulated (B) ccsRNAs between NB and ND conditions.

### Characterization of primary csRNA transcripts

Precise end-mapping of the primary transcript was performed by circular RT-PCR. The ends of three primary csRNA transcripts (Cre-csRNA_61, Cre-csRNA_62, and Cre-csRNA_91) were successfully determined. The primary lengths of the csRNAs are as follow: Cre-csRNA_61 (271 nt), Cre-csRNA_62 (220 nt) and Cre-csRNA_91 (165 nt) as shown in Table S4. These three primary transcripts were subsequently subjected to a secondary structure prediction. All the primary csRNA transcripts formed multiple stem-loop structures (Figure 4). When a BLAST analysis was performed on all the primary csRNA transcript sequences using a cut-off E-value of 1 x 10^-4^, only microalgae chloroplast genomes were found to have the highest identity values as shown in Table 5. Interestingly, 46% region of csRNA_91 primary transcript was shown to be 78% identical with the genome sequence of a cyanobacterium of *Rivularia sp*. PCC 7116.

**Figure 4:**
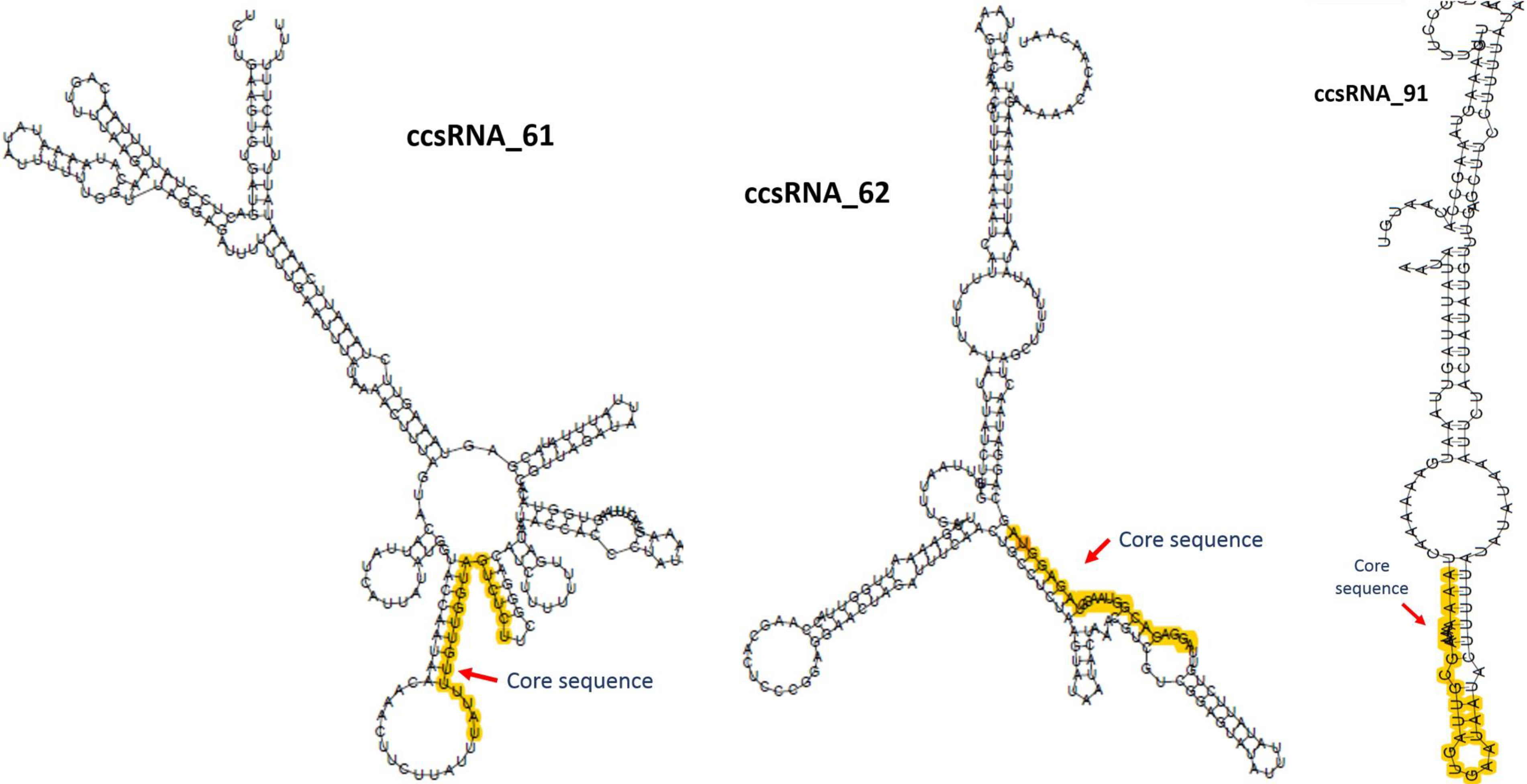
Optimal secondary structure prediction of csRNA primary transcripts using *RNAfold* web server. The core sequences are highlighted in yellow color.

**Table 5.** The interspecies comparison of Cre-csRNA primary transcript

| ccsRNA | Description of BLAST hits | Coverage | E value | Identity |
| --- | --- | --- | --- | --- |
| Cre-csRNA_61 | *Pleodorina starrii* plastid, complete genome | 11% | 7e-05 | 100% |
|  | *Volvox carteri f. nagariensis* strain UTEX 2908 plastid, partial genome | 12% | 7e-05 | 97% |
|  | *Gonium pectorale* chloroplast DNA, complete genome, isolate K3-F3-4 | 11% | 9e-04 | 100% |
| Cre-csRNA_62 | *Tetrabaena socialis* chloroplast, partial genome | 26% | 6e-10 | 98% |
| Cre-csRNA_91 | *Tetrabaena socialis* chloroplast, partial genome | 26% | 6e-10 | 98% |
|  | *Gonium pectorale* chloroplast DNA, complete genome, isolate K3-F3-4 | 29% | 1e-10 | 96% |
|  | *Volvox carteri f. nagariensis* strain UTEX 2908 plastid, partial genome | 43% | 1e-04 | 79% |
|  | *Rivularia* sp. PCC 7116, complete genome | 46% | 1e-05 | 78% |

## DISCUSSION

Non-coding RNAs have emerged as important factors in the regulation of a wide range of biological processes. Besides being encoded in the nuclear chromosomes, they can also be derived from the chloroplast genome and these are known as chloroplast small RNAs (csRNAs). However, information on the functionality of csRNAs is limited. A comparative approach by using different growth conditions may provide hints on their roles. N deprivation has been shown to adversely affect chloroplast functions in the model unicellular microalga *C. reinhardtii* (Plumley and Schmidt 1989; Juergens *et al*. 2015) In this study, the differential accumulation of csRNAs in *C. reinhardtii* was examined under N deprivation and their potential roles were investigated. The locations of differentially accumulated csRNAs were mapped, and their potential mechanisms of actions are postulated on the basis of their positions.

Early studies on plant csRNAs focussed more on the profiling of stress-responsive genes by identification. Subsequently, the differential accumulation of csRNAs under heat stress were observed in Chinese cabbage (Wang *et al*. 2011). A similar study was also performed in barley to understand the differential accumulation of csRNAs during drought stress and phosphorus-deficiency (Hackenberg *et al*. 2013; Hackenberg *et al*. 2015). Nevertheless, there is yet an attempt to link csRNAs to N availability stress. This physiological condition offers a good prospect of uncovering csRNAs that are differentially expressed as well as their possible roles. Such studies were mostly done on nuclear-encoded microRNAs (miRNAs). In *C. reinhardtii*, the expression patterns of miRNAs have been shown to be affected by individual deprivation of either N (Voshall *et al*. 2017), sulfur (Shu and Hu 2012) or phosphate (Chaves Montes *et al*. 2014). In terms of function, miRNAs have been shown to regulate target protein-coding genes by either translational inhibition or mRNA cleavage (Chung *et al*. 2017; Lou *et al*. 2018; Wang *et al*. 2018). Unfortunately, csRNAs were not given due attention in these previous studies. This led to our search for differentially accumulated csRNAs during nutrient stress, specifically N deprivation, and their potential physiological roles.

### Effect of N deprivation on the abundance of Cre-csRNAs

This study demonstrated the differential level of various csRNAs during N deprivation. Our understanding of the biogenesis of csRNAs is still limited and perhaps the determination of their genomic locations and the neighboring genes might be useful. Mapping of the genomic locations of the differentially accumulated csRNAs revealed that they originated mostly from either untranslated regions or sequences coding for ends of tRNA and rRNA. They could also be derived from intergenic regions or generated in the antisense direction.

The original positions of the csRNAs coding regions could also provide a deeper understanding of their potential binding sites and mode of action. Among those that were mapped to the intergenic regions, 6 differentially accumulated csRNAs were found to be located at the 5’ untranslated transcribed regions (UTRs) of protein-coding genes. This class of csRNAs are found a short distance upstream of start codons. It was discovered that the sequence of these csRNAs coincided with the 5’ termini of chloroplast mRNAs. It is at present unclear, whether any of these csRNAs serve a specific function. As described in previous literature, these csRNAs might involve in mRNA stability by serving as a decoy. An accumulation of csRNA serves as the binding sites of pentatricopeptide repeat proteins (PPR). PPRs, bound to the target RNA, guide the exonuclease to target and degrade the latter (Ruwe and Schmitz-Linneweber 2012). This action protects mRNAs that have the same csRNA sequence from exonucleolytic degradation by shifting the exonuclease towards the actual csRNA molecules. Hence, the amount of csRNAs were purposefully accumulated to avoid the mRNA of the targeted genes from degradation.

Accumulation of csRNA can also lead to a reduction in the expression of its target gene could also be possible. This inverse relationship showed that an alternative mechanism different from the PPR-facilitated mode of action is available. This outcome is usually shown by the classical gene silencing mechanism in which the upregulation of small RNAs leads to the downregulation of their target genes. In gene silencing, small RNAs guide the cleavage proteins such as Argonautes to the specific target sequence by sequence complementarity. This leads to cleavage of the targeted mRNAs resulting in translational inhibition (O’Brien *et al*. 2018). This combination of small RNAs and proteins is known as RNA-Induced Silencing Complex (RISC). The mechanism is widely studied in miRNAs of various organisms including *C. reinhardtii* (Zhao *et al*. 2007; Cerutti *et al*. 2011; Chung *et al*. 2017; Wang *et al*. 2018). Interestingly, mitochondria possess such a miRNA-mediated gene silencing mechanism and the miRNA molecules are encoded within the mitochondria genome (Barrey *et al*. 2011; Latronico and Condorelli 2012). On the other hand, such a circumstance does not occur in the chloroplast and our *in silico* search for chloroplast-encoded miRNA candidates was unsuccessful (data not shown). Genes coding for RISC proteins were also not found in the chloroplast genome. In addition, none of the nuclear-encoded RISC proteins have chloroplast signal peptide. Thus far, there has been no report on the exportation of nuclear-encoded RISC proteins into the chloroplast. All these circumstances pointed towards an absence of a miRNA mediated gene silencing phenomenon inside the chloroplast. Therefore, an opposite correlation between these csRNAs and their targeted genes potentially opens up the possibility of a new silencing mechanism in the chloroplast and this requires further investigation.

In the case of the csRNAs located at 3’-UTR of protein-coding genes, four differentially accumulated csRNAs were found and they were all upregulated. The sequences of these csRNAs are identical to the sequences of the 3′-UTR of protein-coding genes. These csRNA molecules are potentially produced by either transcription from internal promoters or processing of the associated mRNAs. The mode of action of this type of csRNA is still unknown. In prokaryotes, small RNAs originating from 3’-UTR of protein-coding genes have been observed and their mechanism of action elucidated. The 3’-UTR end of cpxP mRNA was cleaved by the enzyme RNase E to release a small RNA called CpxQ (Miyakoshi *et al*. 2015). CpxQ, together with the Hfq protein, represses mRNAs of envelope proteins, a pathway that protects the bacteria against any inner membrane damage (Chao and Vogel 2016). From our study, the upregulated csRNAs are located at the 3’-UTR of the protein-coding genes *atpA*. *psbI*, *psbF*, and *psaJ*. All of these genes are involved in light absorption and electron transfer during photosynthesis. Differential accumulation of these csRNAs during varying N availability showed their involvement in the regulation of photosynthesis. It remains unknown whether they act on the mRNAs they originated from or others. As the Hfq-mediated mechanism can only be found in prokaryotes, the actual mode of action of the *Chlamydomonas* csRNAs requires further investigation.

Intergenic regions can also serve as generators of noncoding RNAs (ref). Both strands of the chloroplast genome were found to be fully transcribed (Shi *et al*. 2016). This led to the transcription of all regions enabling the production of short RNA molecules after post-transcriptional processing (Hotto *et al*. 2012). Despite a high percentage of span in the chloroplast genome, the number of differentially N-responsive csRNAs originating from this region was low. Only one was found to be upregulated and another was downregulated. The rest of the short RNA molecules might not play any significant role in N availability. They could be possibly differentially accumulated under different physiological conditions.

The most abundant csRNAs were found in the ends of chloroplast rRNA and tRNA genes. The DNA sequences representing these csRNAs overlapped with the terminal sequences of the rRNA and tRNA genes. They could be post-transcriptional cleavage products of these genes. The rRNA-derived csRNAs were preferentially located at the 3’-ends of the rRNAs, while the tRNA-derived csRNAs were mainly located at 5’-terminals of tRNAs. A study in Chinese cabbage was the first to identify both types of csRNAs and showed that their accumulation was responsive to heat stress. The abundance of these csRNAs increased as the heat stress was introduced (Wang et. al., 2011). The similar csRNAs were found to be differentially accumulated in phosphorus-deprived barley, but they were all downregulated. In the case of N deprivation on *Chlamydomonas* cells, only six csRNAs from these regions were found to be differentially accumulated and all were downregulated. Here, it was demonstrated that abiotic stress affects the accumulation of these csRNAs, which might demonstrate their importance. These csRNAs could possibly play a role in regulating the stability of tRNAs and rRNAs during stress. On the other hand, a study in *Arabidopsis thaliana* found that these csRNAs accumulate outside the organelle and this led to a hypothesis that they could serve as a signaling pathway between the nucleus and chloroplast (Cognat *et al*. 2017).

Antisense csRNAs were mostly reported in the findings of noncoding RNAs in chloroplast. In this study, three csRNAs were antisense to the gene *atpB* were differentially accumulated and they were all upregulated. A possible role of this type of csRNAs in *atpB* regulation was previously described. A chloroplast genome rearrangement in *C. reinhardtii* upregulated the expression of a csRNA that was antisense to *atpB*. This csRNA restabilized a previously unstable *atpB* mRNA which had a deleted 3’ stem-loop segment (Hotto *et al*. 2010). Besides, two downregulated csRNAs were antisense to the *rbcL* gene and they were all downregulated. A positive correlation was also observed for antisense csRNAs and their putative mRNA targets (Chen *et al*. 2014). However, a further investigation is needed to ascertain the mechanism of action of this type of csRNAs on their target genes.

Understanding the molecular mechanisms of each type of csRNAs is important to distinguish their mode of action. Previous studies have successfully revealed the action of small RNAs by elucidating their structures as well as recognizing the proteins they associate with (Kim *et al*. 2010). Proteins play a vital function in the action of small RNAs. These include the actions of Argonaute and DICER proteins with microRNAs (Hammond 2005), Cas9 with CRISPR RNAs (Jansen *et al*. 2002), Piwi with piRNAs (Seto *et al*. 2007), and Hfq with bacterial small RNAs (Vogel and Luisi 2011). Therefore, finding the associated proteins is important to elucidate their mechanism. A few RNA-binding proteins have been recognized in the chloroplast of *Chlamydomonas*. They were found to be involved in RNA processing by acting as endonuclease and exonuclease (Lurin *et al*. 2004; Schmitz-Linneweber 2008; Pfalz *et al*. 2009) Whether these proteins are actually associated with the csRNAs described in this study remains to be elucidated.

### Cre-csRNAs are conserved in microalgae

The primary sequence of three Cre-csRNAs showed high identity with sequences within the microalgal chloroplast genomes. However, there is only a limited number of complete algal chloroplast genomes available in the current database. None of the hits included the chloroplast genomes of plants indicating the possibility that these Cre-csRNAs are phylogenetically conserved only among the microalgae. Interestingly, a stretch of sequence within the Cre-csRNA_91 primary transcript matched the genome sequence of the cyanobacterium *Rivularia* sp. PCC 7116. Assuming that cyanobacteria are the ancestors of present-day chloroplasts, similar noncoding RNAs might still be present in some cyanobacterial species and possibly play the same physiological function (Raven and Allen 2003; Kopf and Hess 2015).

### Potential regulatory function of csRNAs during N deprivation

Nitrogen is involved in the physiological changes in the chloroplasts and its availability evidently affects the expression of csRNAs. A previous study on *C. reinhardtii* demonstrated that N deficiency decreased the rate of synthesis of photosynthetic proteins, especially the Light-Harvesting Complex apoproteins (Plumley and Schmidt 1989). The levels of transcripts and proteins of photosystems I and II were also downregulated under N deprivation (Juergens *et al*. 2015). A possible mode of regulation of chloroplast protein synthesis is by inhibition of translation. In this study, several differentially accumulated csRNAs were found during N deprivation of *C. reinhardtii* and some of these csRNAs are possibly involved in posttranscriptional processing of chloroplast transcripts. Further studies are needed to determine the presence of this type of regulation. Nevertheless, these findings provided new insights into the involvement of non-coding RNAs in chloroplast RNA metabolism during N deprivation.

## CONCLUSION

In this study, we demonstrated that N deprivation affected the level of csRNAs in *Chlamydomonas reinhardtii*. The affected csRNAs originated from different genomic locations that represent either untranslated, intergenic, antisense or ends of tRNA and rRNA. All csRNAs derived from 3’-UTR regions were upregulated while all csRNAs derived from the ends of tRNA and rRNA were downregulated. The regulations of these csRNAs indicated that they have physiological roles during N deprivation. Their importance is reflected by the high sequence identity of three primary csRNA sequences with the chloroplast genomes of a few microalgal species.

## ACKNOWLEDGMENTS

This work was supported by the Malaysian Ministry of Education under the Fundamental Research Grant Scheme (Grant No. 203/PBIOLOGI/6711458). The authors have no conflict of interest to declare.

## AUTHOR CONTRIBUTIONS

S.C. was responsible for conception and design, data collection, results analysis and manuscript writing; G.A., and N.N. for conception and design, data analysis, critical revision and final approval of the manuscript. All authors read and approved the final manuscript.

